# The interplay of spatial scale and landscape transformation moderates the abundance and intraspecific variation in the ecomorphological traits of a phyllostomid bat

**DOI:** 10.1101/2021.05.01.442263

**Authors:** Andrés F. Ramírez-Mejía, J. Nicolás Urbina-Cardona, Francisco Sánchez

**Author notes:** Corresponding author. Instituto de Ecología Regional, UNT – CONICET. Residencias Universitarias de Horcomolle, Edificio las Cupulas. ZIP 4107.

## Abstract

Land-use intensification imposes selective pressures that systematically change the frequency of wild population phenotypes. Growing evidence is biased towards the comparison of populations from discrete categories of land uses, ignoring the role of landscape emerging properties on the phenotype selection of wild fauna. Across the largest urban-rural gradient of the Colombian Orinoquia, we measured ecomorphological traits of 216 individuals of the Flat-faced Fruit-eating Bat *Artibeus planirostris*, to evaluate the scale of effect at which landscape transformation better predicts changes in phenotype and abundance of an urban-tolerant species. Forest percentage at 1.25 km was the main predictor affecting abundance, wing aspect ratio, and body mass of this phyllostomid; but the direction of the effect differed between abundance and ecomorphological traits. Although landscape factors explained changes in the forearm length at all spatial scales, the effect was sex-dependent and the most important predictor was forest percentage at 0.5 km. Our results indicate that landscape elements and spatial scale interact to shape ecomorphological traits and the abundance of *A. planirostris*. Interestingly, the scale of effect was congruent among all biological responses. A pattern that likely arises since species’ abundance can reflect the variation on phenotype under different environmental filters across landscape scenarios.

## INTRODUCTION

Land-use intensification imposes selection regimens that influence the composition and structure of biotic communities (Echeverría-Londoño et al. 2016). Species that are able to adapt—or even prosper—in anthropogenic landscapes, are subject to environmental filters that lead to changes in the frequency of phenotypes across populations (Alberti et al. 2017). That is, human environments might promote directional selection processes that alter the ecomorphological traits of the species (e.g., Puckett et al. 2020). This pattern has been increasingly documented in several vertebrates such as lizards (Lazić et al. 2013, Hall & Warner 2017, Putman & Tippie 2020), small non-volant mammals (Snell-Rood & Wick 2013, Puckett et al. 2020), bats (Tomassini et al. 2014), birds (Liker et al. 2008, Caizergues et al. 2021), and middle-sized carnivores (Parsons et al. 2020). Nevertheless, most studies focused on comparing populations between discrete categories of land-use intensification, e.g., urban vs. rural, ignoring the role of landscape emergent properties and the spatial scales that better correlate with phenotypic changes of fauna in human environments (Strubbe et al. 2020).

New World bats of the family Phyllostomidae constitute an ideal model to study landscape transformation effects on species’ phenotype, since it includes several species that are tolerant to habitat transformation (Meyer et al. 2016, Farneda et al. 2020), and they are highly variable in ecomorphological traits that drive their adaptation to environmental change (Soriano 2000, Farneda et al. 2015). Physical constraints on phyllostomids’ wing morphology and body size impose aerodynamical restrictions to exploit open or cluttered environments (Farneda et al. 2015, Wordley et al. 2017). That is, phyllostomid species with low wing aspect ratio and low wing loading are adapted to slow and maneuverable flight in structurally complex habitats with dense vegetation (Marinello & Bernard 2014). Consequently, they are expected to be highly sensitive to habitat loss and disruption. Alternatively, bats with high wing aspect ratio and high wing loading can fly at a high speed and are aerodynamically efficient in open areas (Norberg & Rayner 1987, Marinello & Bernard 2014). Thus, they are thought to be better adapted to survive in deforested landscapes (Wordley et al. 2017). These theoretical predictions have been empirically corroborated in several Neotropical locations (e.g., Meyer & Kalko 2008, Jung & Kalko 2010, Farneda et al. 2015, Ramírez-Mejía et al. 2020). However, most studies focused on the phyllostomid trait responses at the assemblage or trophic guild level (Gonçalves et al. 2017, Brändel et al. 2020, Farneda et al. 2020), overlooking the potential intraspecific consequences of landscape transformation on bats’ ecomorphological traits.

Individuals within a phyllostomid population maximize the use of the space by adapting intraspecific foraging movements (see Kerches-Rogeri et al. 2020). Therefore, bats of a population inhabiting a spatially heterogeneous landscape, such as those created by humans, might face different environmental pressures on their ecomorphological traits that determine their space use compared to those living in wild environments. Few works have attempted to explore the relation between intraspecific ecomorphological traits of phyllostomids with environmental gradients at large geographical scales (see Marchán-Rivadeneira et al. 2012, Stevens et al. 2016), and none have explicitly incorporated landscape effects such as habitat amount or deforestation. Thus, we studied how landscape transformation and spatial scale moderate the intraspecific variation of phyllostomids’ ecomorphological traits. We used as a model the Flat-faced Fruit-eating Bat, *Artibeus planirostris* (Spix 1823), a common phyllostomid species in human-altered environments in the Neotropics (Jara-Servín et al. 2016, Saldaña-Vázquez & Schondube 2016). *A. planirostris* is a relatively large bat, weighing 40 – 69 g (see Hollis 2005), feeding in the canopy on fruiting trees with asynchronous, massive, and short duration production, e.g., *Ficus* spp. (Soriano 2000). This bat regularly roosts in the foliage and changes shelter according to their foraging movements; thus, its home range is expected to be large and variable (Soriano 2000, Hollis 2005).

In this work, we tested the hypothesis that forest replacement leads to the environmental filtering of inter-individual variation in bats’ phenotypes based on their flight performance. Thereto, we predict that non-forested areas, such as anthropogenic grasslands and urbanizations, will positively affect the average body mass and wing traits of *A. planirostris*, since those traits correlate with the bats’ ability to exploit open environments (Jung & Threlfall 2016, Meyer et al. 2016). In addition, since large *Artibeus* species tend to prosper in Neotropical anthropogenic areas (Saldaña-Vázquez & Schondube 2016), we expect that the abundance of *A. planirostris* will be positively affected by forest loss. Nightly foraging movements of *A. lituratus* — a close relative of, and morphologically similar species to, *A. planirostris*—in deforested areas range from ~600 to ~2000 m (1158.8 ± 598.6 m. See Table 1 in Trevelin et al. 2013). Accordingly, we expect *A. planirostris*’ abundance to be affected by landscape transformation at either local (i.e., 500 m), or landscape scales (i.e., 2000 m). However, since the average foraging movement of *A. planirostris* is estimated at around 1000 m (Trevelin et al. 2013), we expect that the landscape effect will be particularly strong at this intermediate scale of our analysis. Finally, since the frequency of phenotypes across a population is not evenly distributed, i.e., some phenotypes are more abundant than others (Bolnick et al. 2003), we expect that the spatial scale which best predicts changes in *A. planirostris* abundance, will be congruent with the spatial scale which shows the best correlation with changes in ecomorphological traits.

**Table 1.**
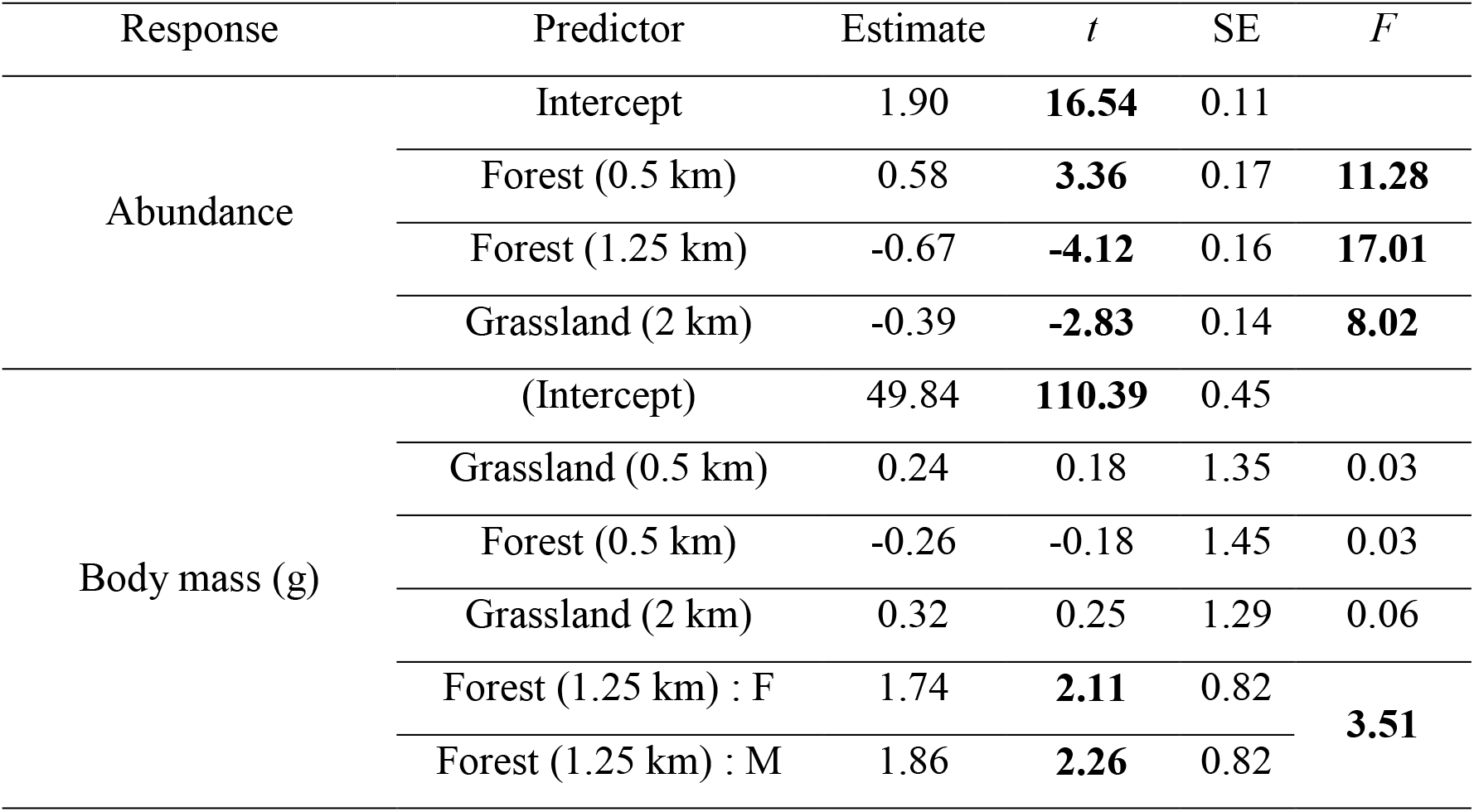
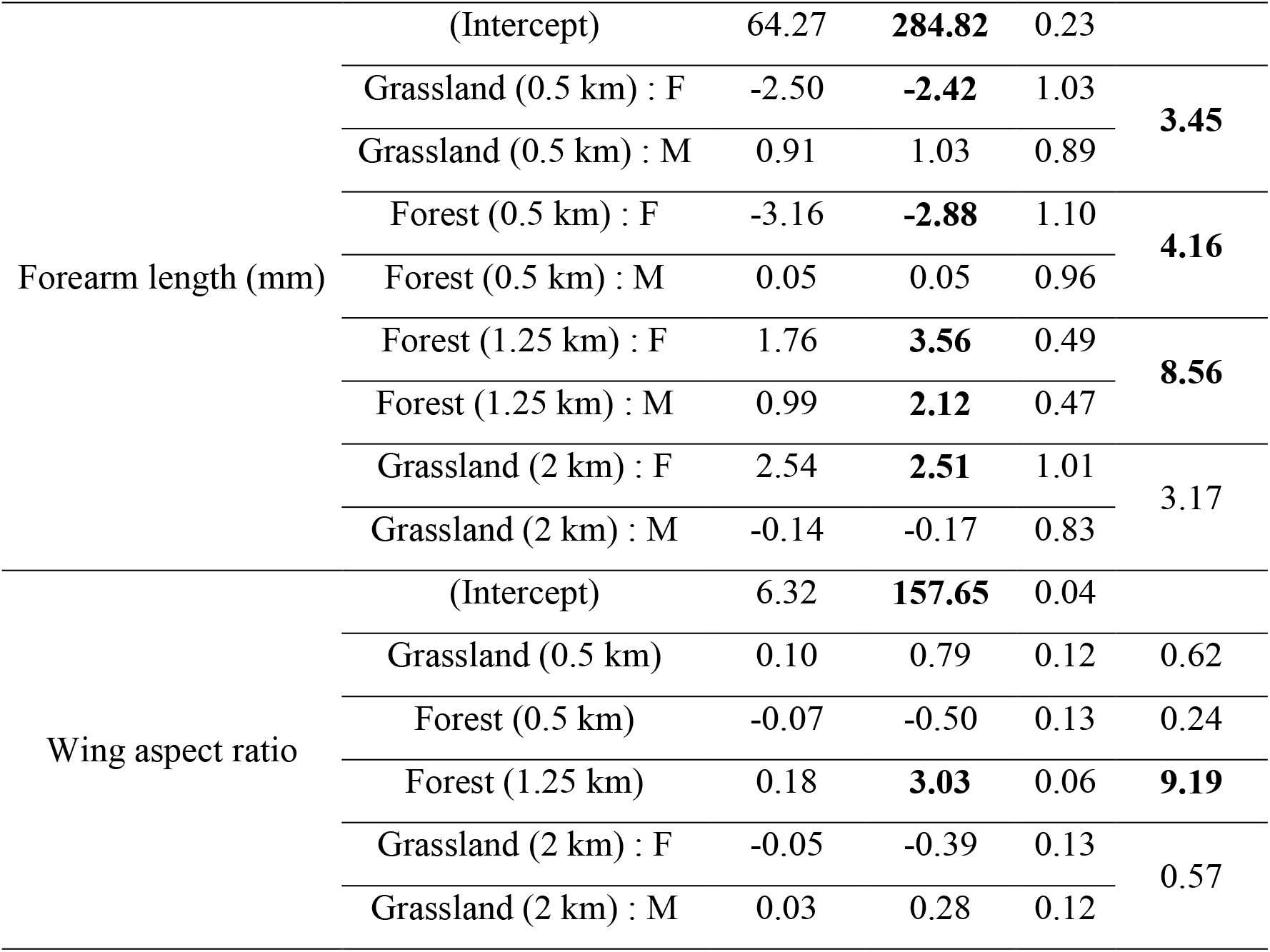
Best-fitted generalized linear mixed effect models explaining changes in *Artibeus planirostris* abundance and its ecomorphological traits in an urban-rural landscape of the Colombian Orinoquia. Bold numbers of *t*-statistics denote that the parameter is significantly different from zero. Bold numbers of *F*-statistics denote that the parameter significantly (p<0.05) reduced the model’s deviance.

## METHODS

### Study area

We tested our hypotheses on an urban-rural gradient on the Andean piedmont of the Llanos Orientales of Colombia, the most densely populated and urbanized zone in the Colombian Orinoco (Villavicencio municipality). The Andes piedmont is a transition area between the Orinoco savannahs and Andean ecosystems (Rangel et al. 1997), and it has experienced profound anthropogenic land cover changes since the early 1950s (Etter et al. 2008, Correa Ayram et al. 2020). Currently, the landscape of this region is a mosaic of a continuous urban core, scattered suburban settlements, forest remnants, and agricultural systems embedded in an extensive anthropogenic grassland matrix (Romero-Ruiz et al. 2012, Sánchez-Cuervo et al. 2012). The rainy season occurs between April and October, whereas the low precipitation season takes place between November and March with an annual mean rainfall of ~2,600 mm (Bernal et al. 2013). The natural vegetation has been described as a rainforest with a strong influence of the Andean piedmont (Bates 1948, Blydenstein 1967, Rangel et al. 1997).

### Bat sampling

Along Villavicencio’s urban-rural gradient, we made a non-random selection of five sites to maximize the variability of predictor variables while reducing the spatial autocorrelation among landscape units (LU) (Figure 1). Hereafter, LU refers to a set of concentric circles in which we measured landscape composition and structure; and sites refer to the sampling units set in the centroid of each LU (Figure 1). We chose sites that had enough tree cover to place mist-nets to capture bats (the canopy cover above mist-nets at all sites was >80%), and a linear separation of at least 4.3 km from the nearest sampling site to avoid LUs overlap (Figure 1). We visited each site five times between January and July of 2016, setting two to three 12×3m mist-nets during nights with moon illumination levels below 60% (Rowse et al. 2016). We used a rabbit tattoo to individually mark each bat with a unique numerical code (Powell and Proulx 2003) to guarantee the independence of bat captures. We placed the tattoo in the lower part of the plagiopatagium of the right wing. Sampling effort at each site ranged from 288 to 300 m^2^×nights, and the total sampling effort was 1,484 m^2^×nights.

**Figure 1.**
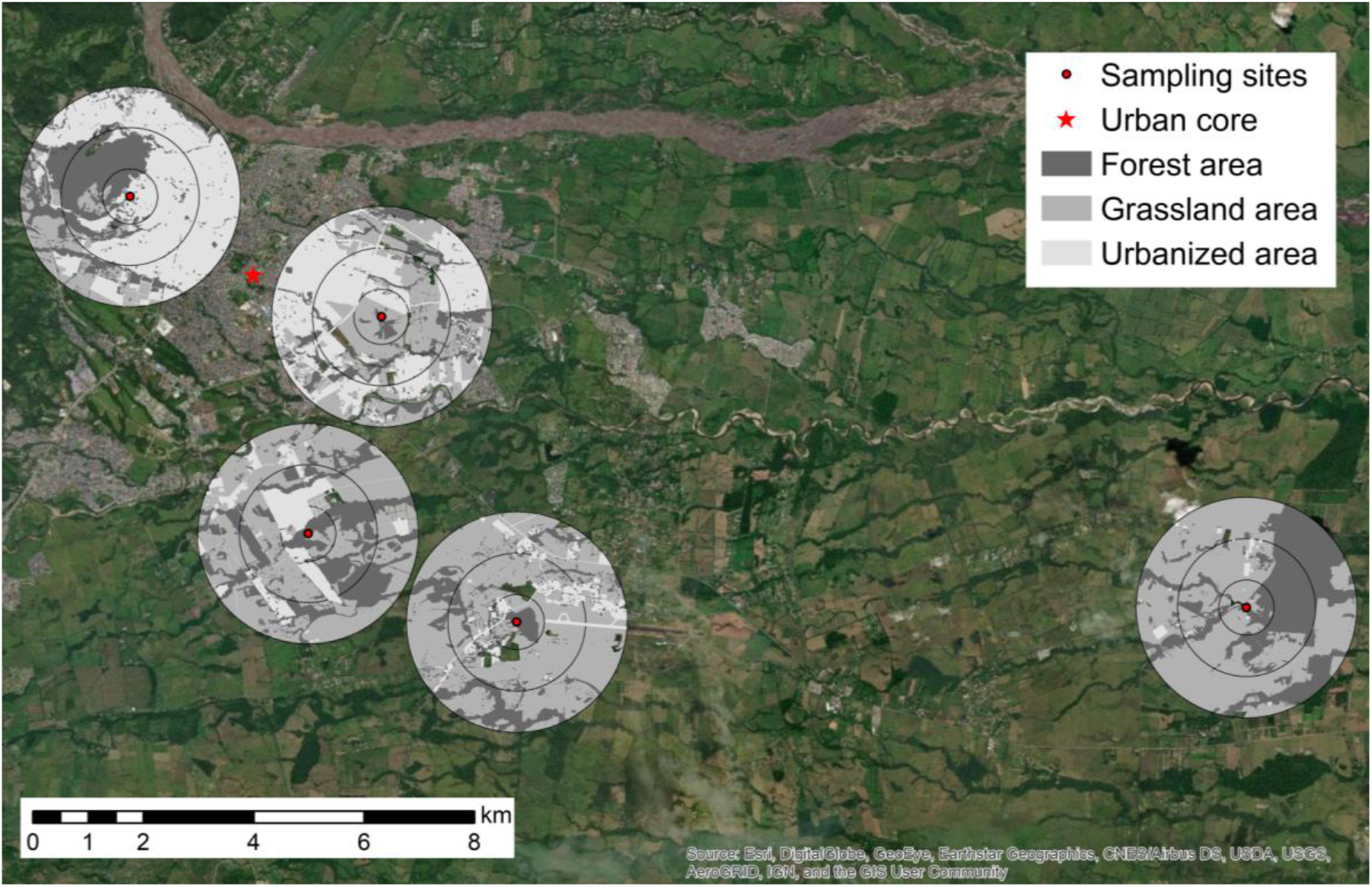
Landscape units (LU) in an urban-rural landscape in Villavicencio, Colombian Orinoco basin. We sampled bats at the centroid of each concentric circle (sampling sites). Concentric circles at each LU set the limits of each spatial scale (0.5, 1.25, and 2 km of radius) in which landscape variables were measured.

### Ecomorphological traits

We considered four morphological traits related to the bats flight performance: wing loading, wing aspect ratio, body mass, and forearm length. Wing loading and wing aspect ratio are mainly related to flight speed and maneuverability (Marinello & Bernard 2014), whereas forearm length and body mass are mainly associated with phyllostomid flight speed (Morrison 1980, Winter 1999). We used a 0.01 mm resolution caliper to measure forearm length following Díaz et al. (2016), and measured body mass using a digital balance of 0.1 g resolution. Wing loading and wing aspect ratio were measured from a photograph of the left wing fully extended in a ventral position against a paper with millimeter scale (Korine & Pinshow 2004). These images were processed using ImageJ 1.6 (Abramoff et al. 2004) to measure wing span and wing area as described by Norberg & Rayner (1987). Finally, we calculated the wing aspect ratio as the quotient of the wing span squared and wing area, and estimated wing loading by dividing the body mass into the product of gravitational acceleration, 9.81 m/s^2^, and wing area (Norberg & Rayner 1987). We only measured morphological traits of adults and non-pregnant females.

### Land use classification

We delimited concentric buffers of 0.5, 1.25, and 2 km (local, intermediate, and large spatial scales, respectively) of radius around the sampling sites to characterize landscape composition and structure (see Miguet et al., 2016). These spatial scales were expected to encompass the minimum, maximum, and average foraging movements reported for other large *Artibeus* species (see Trevelin et al., 2013). We performed a classification on ArcGIS 10.1, using DigitalGlobe satellite images taken between 2014 and 2015 with a 2 m^2^ per pixel resolution (Google Earth 2016), to generate maps with three land uses: built area, anthropogenic grassland, and native forest (Figure 1). At each spatial scale, we measured the percentage of built-up areas, grassland, and forest cover, the number of forest patches, mean forest patch size, forest patch size standard deviation, and density of forest patches (Mendes et al., 2016; Pinto & Keitt 2008). We did not consider other land uses and landscape elements (e.g., crops, water bodies, or bare ground) in the analysis, because they covered less than 5% of each area at a 2 km radius, and they were not present in each LU’s (Figure 1).

### Data analyses

We performed Pearson correlations among landscape predictors and morphological response variables to identify linear relationships and excluded highly correlated predictor variables from further analyses (*r* > 0.7) (Quinn & Keough 2002). We used generalized linear mixed models (GLMM) to analyze changes in abundance and ecomorphological traits of *A. planirostris* as a function of landscape metrics. The first model included abundance data of *A. planirostris* as the response variable, i.e., the number of captures by site, and the landscape variables at different scales as predictor variables. In subsequent models, we used as responses variables the average value of each ecomorphological trait by sex, from the individuals captured in each landscape unit. As predictors, we used the landscape variables at several spatial scales and fixed the bats sex as an interaction term of all predictor variables to account for sexual dimorphism (Figure 2). We included the log-transformed number of captured individuals of each sex as an offset variable in these models, to consider the potential effect of the sample size on the estimation of the average value of morphological traits (see Zuur et al. 2009). In all models, we used the LU as a random term to account for potential spatial autocorrelation issues (Rhodes et al. 2009). We evaluated the fit of *A. planirostris* abundance and morphological traits to normal, log-normal, gamma, Poisson or negative binomial distributions—according to the case—using the package *fitdistrplus* of R 3.6.3 (Delignette-Muller & Dutang 2015, R Core Team 2020). We used z-scores of non-correlated predictor variables to compare parameter importance in the model (Marquardt 1980), and fitted all models using the *lme4* package (Bates et al. 2015). We estimated p-values of models’ parameters via Satterthwaite’s degrees of freedom method with the *lmerTest* package (Kuznetsova et al. 2017) and calculated the conditional and marginal R^2^ of each model using the *MuMin* package (Bartón 2016). Finally, for model selection, we applied a likelihood ratio test (Anisimova & Gascuel 2006), and we validated the model fit using graphical analysis of residuals. We did data wrangling and plotted partial residuals of best-fitted models using R’s base functions and the packages *dplyr, cowplot*, and *ggplot2* (Wickham 2016, Breheny & Burchett 2017, Wilke 2019, Wickham et al. 2020).

**Figure 2.**
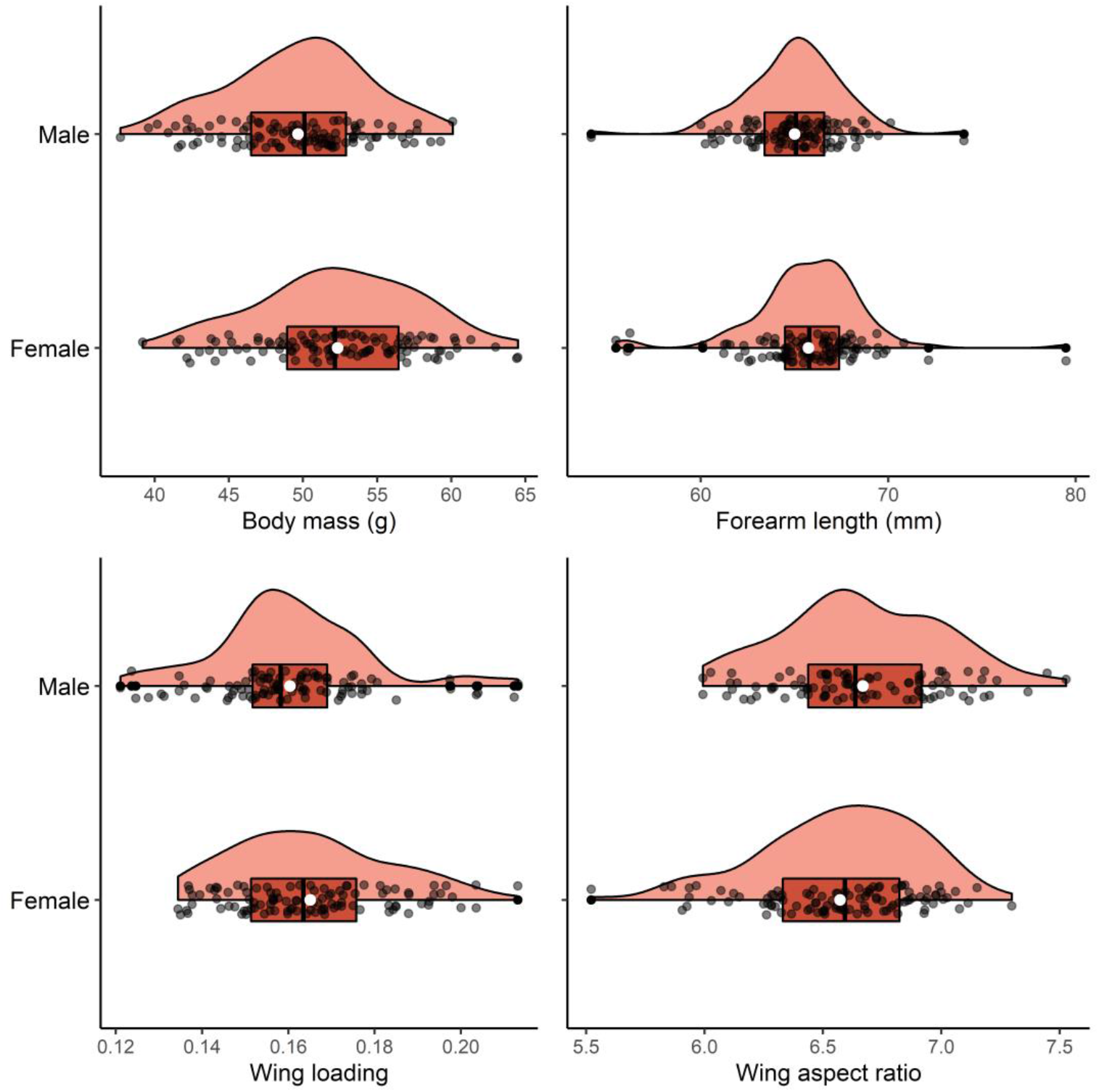
Boxplots and density plots of *Artibeus planirostris* ecomorphological traits in an urban-rural landscape of the Colombian Orinoco basin. Black dotes: observations. White dotes: average value of the trait.

## RESULTS

We captured 232 individuals of *A. planirostris* and measured the ecomorphological traits of 216 adult males and non-pregnant females (Figure 2). During the sampling, we recorded two recaptures in two LU, and both times, recaptured bats were found at their original LU. For additional details regarding sampling and the phyllostomid assemblage recorded see Ramírez-Mejía et al. (2020). After correlation analyses, we excluded landscape collinear variables and retained grass cover at 0.5 and 2 km scales and forest percentage at 0.5 and 1.25 km (Figure S1). All response variables fitted a normal or log-normal distribution (link function = identity), and the correlation coefficients of morphological response variables were below 0.5 (Figure S2). The abundance of *A. planirostris* was explained by landscape predictors at all spatial scales (Figure 3). The percentage of forest at 1.25 km negatively affected the *A. planirostris* abundance and was the most important variable in the model, followed by forest at 0.5 km and grassland at 2 km (Figure 3, Table 1). The model explained 42.58% of the *A. planirostris* abundance (42.58% marginal R^2^ and 42.58% conditional R^2^). Residual analysis showed that the model was well fitted (Figure S3).

**Figure 3.**
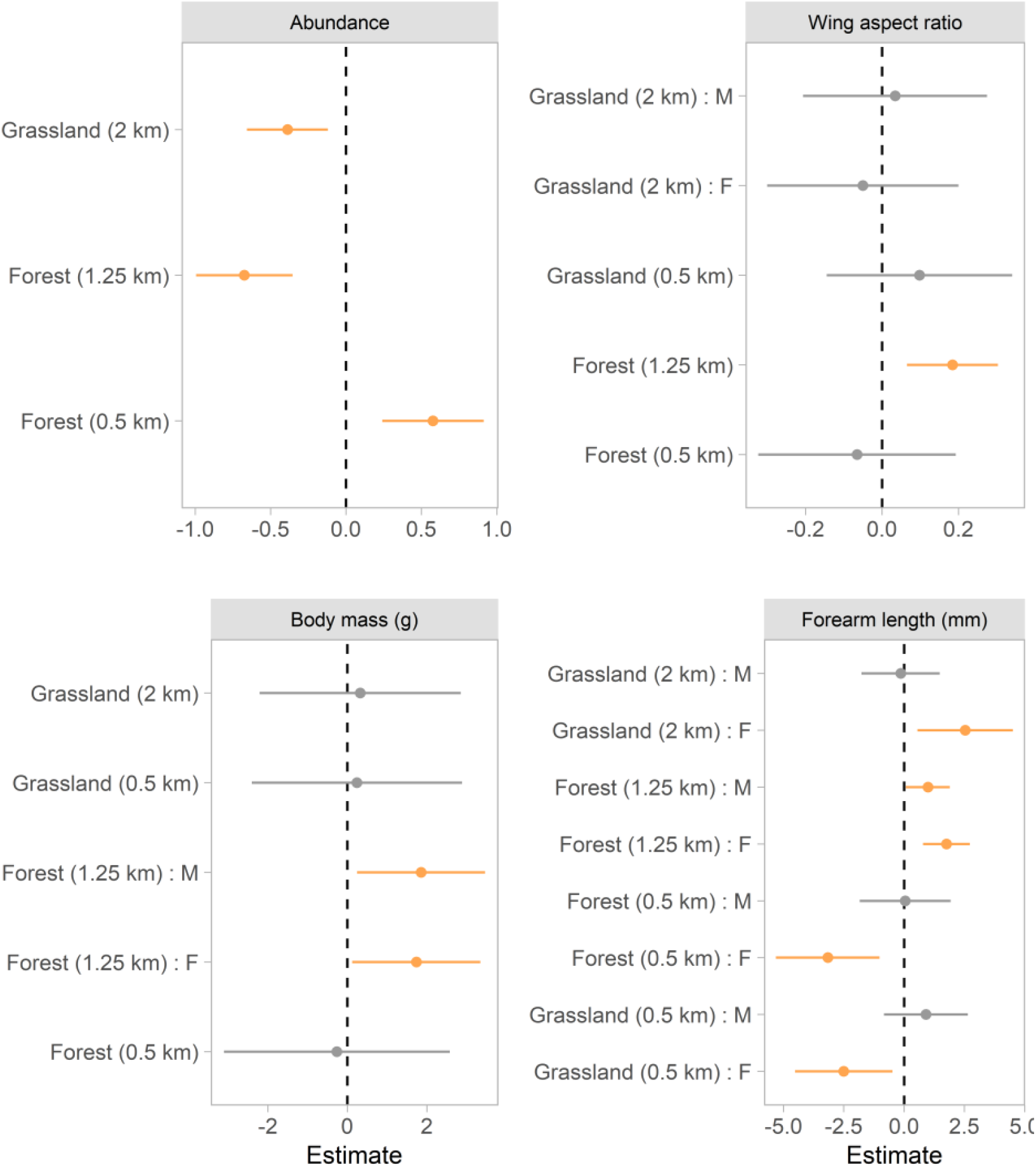
Best-fitted generalized linear mixed effect models explaining changes in *Artibeus planirostris* abundance and ecomorphological traits, in an urban-rural landscape of the Colombian Orinoco basin. Orange parameter (dots) and confidence intervals (horizontal lines) indicate that the slope is statistically different from zero and the parameter significantly reduced the model’s deviance. M: male. F: female.

The best-fitted model explaining wing aspect ratio variation of *A. planirostris* included grassland at 0.5 and 2 km, and forest at 0.5 and 1.25 km scales; however, only forest percentage at 1.25 km had a significant positive effect (Figure 3). This model explained 27.65% of the total variation (27.65% marginal R^2^ and 27.65% conditional R^2^). Forest percentage at the 1.25 km scale also explained changes in body mass (Figure 2). This predictor had a positive effect and depended on bats’ sex (Figure 3). The model also included non-significant predictors and explained 20.82% of the total variation (20.82% marginal R^2^ and 20.82% conditional R^2^). Forearm length of *A. planirostris* was significantly affected by landscape variables at all spatial scales and showed a sex-dependent response as well (Figure 3, Table 1). The forearm length was positive associated with forest percentage at 1.25 km, but the effect tended to be higher for females (Figure 3). Forest and grassland percentages at 0.5 km negatively affected the forearm length of females, whilst it did not show a significant relation for males (Figure 2). Similarly, the grass percentage at 2 km had a positive effect on females’ forearm length, but no effect for males (Figure 2). Forest percentage at 0.5 km was the most important variable in the model, followed by the grass percentage at 0.5 and 2 km scales, and the forest percentage at 1.25 km (Table 1). This model explained 32.76% of the total variation (32.76% marginal R^2^ and 32.76% conditional R^2^). All models of ecomorphological traits exhibited a good fit (Figures S4 – S6), except for wing loading and we did not further consider it.

## Discussion

Our results support the hypothesis that forest replacement leads to environmental filtering of inter-individual variation of bats’ phenotypes, specifically the ecomorphological traits related to flight performance. Indeed, we found that landscape transformation operates at particular spatial scales to affect phenotype and the abundance of an urban-tolerant phyllostomid species. The data support two of the three predictions evaluated in this study. As expected, forest cover at 1.25 km was the most important predictor variable, negatively affecting *A. planirostris* abundance. Body mass, wing aspect ratio, and forearm length responded mainly to forest cover changes at 1.25 km and this was congruent to the abundance response, but contrary to our initial expectation. Remarkably, effects of grassland and forest cover on forearm length varied depending on the spatial scale and sex, reflecting the complexity of landscape effects on the inter-individual phenotype variation in *A. planirostris*.

Our results are in agreement with the general pattern that habitat conversion tends to increase the abundance of generalist phytophagous phyllostomids (Saldaña-Vázquez & Schondube 2016, Gonçalves et al. 2017, Farneda et al. 2020). Bats of the *Artibeus* genus commonly feed on fruits of *Ficus* spp. and *Cecropia* spp. trees (e.g, Saldaña-Vázquez et al. 2013). These plants are common in the anthropogenic matrix of the landscape we studied, and were reported as the main food items of *A. planirostris* in the same area (Ramírez-Mejía 2017). The spatial distribution of these food resources and the nightly foraging movements of the individual bats likely explain the abundance patterns that we found in our study. In anthropogenic landscapes, the average nightly movement of *A. lituratus* is about 1158 m and may range between 418 to 2053 m (Trevelin et al. 2013). Hence, if *A. planirostris* and *A. lituratus* have similar movement capabilities, as expected by their similar ecomorphological traits, this can explain the stronger scale of effect at 1.25 km, since this spatial scale is likely to encompass *A. planirostris* average foraging range (Gorresen et al. 2005, Trevelin et al. 2013). We found a change in the direction of the effect of forest cover on the abundance from the 0.5 km to the 1.25 km scale. This suggests that at local scales *A. planirostris* requires enough forest cover to provide roosting sites and food resources (Mendes et al. 2016), whereas at the larger extent the forest cover may reduce its opportunity to exploit available fruit resources in the deforested matrix (Torres et al. 2018). Besides, the negative relationship with grass cover at the largest scale, 2 km, suggests a detrimental effect of extremely deforested landscapes on the abundance even for long-distance dispersal bats such as *A. planirostris* (Meyer et al. 2016).

Changes in abundance and ecomorphological traits of *A. planirostris* showed consistent relationships with landscape factors and spatial scales. This pattern probably arises since individuals’ abundance can reflect the variation in phenotype prevalence under different selection regimes, in our case landscape scenarios (Bolnick et al. 2003). Indeed, wing aspect ratio, body mass, and forearm length were positively affected by forest cover at 1.25 km regardless of the sex. These results seem counterintuitive — empirical evidence at the community level points to an opposite trend (Jung & Threlfall 2016, Meyer et al. 2016)—and it is likely the result of the particular structure of our studied urban-rural landscape. The forest area in our analysis ranged from 14% to 42% at 1.25 km scale which, in most cases, corresponded to a few large forest patches (see Figure 1). This particular spatial arrangement of natural habitat leads to a large percentage of non-forest area that is probably used by *A. planirostris*, in which case a large body and large-pointed wings may result advantageous (Norberg & Rayner 1987). Indeed, there should be trade-offs that limit the individual specialization of *A. planirostris* to use open environments, without reducing its opportunity to exploit food and roosting resources in structurally complex forests (Bolnick et al. 2003). Thus, the positive relation between forest cover and flight performance traits may mask the underlying complexity of trade-offs regarding bats’ phenotypic response. This interpretation is speculative and deserves further exploration involving trait morphology data, a wider range of forest loss scenarios, and a direct evaluation of how bats differentially use multiple land cover types in anthropogenic landscapes.

Unlike other ecomorphological traits, both forest and grassland cover affected the female’s forearm length at all spatial scales of our analysis. Particularly, forest cover at 0.5 km showed the strongest negative effect. Pregnancy state and the transportation of newborn offspring might impose different aerodynamic challenges for female and males bats (Camargo & Oliveira 2012); that is, during particular reproductive stages, phyllostomids’ flight related-traits might experience different intensity of selective pressures depending on the individual’s sex. Concerning female *A. planirostris* individuals, short forearm lengths should lead to slower and more maneuverable flight, which is advantageous in structural complex forested areas (Morrison 1980, Stockwell 2001). Natural areas are likely fundamental for offspring shelter and access to more diverse food resources during the lactation period (Durant et al. 2013). On the contrary, the positive relation between female forearm lengths and grassland cover, at 2 km scale, could be due to the selection of an aerodynamic efficient flight in non-forest areas. The positive association of bats’ forearm length and flying speed has been extensively documented at the community level (Hayward & Russell 1964, Morrison 1980, Winter 1999), and, similarly, might be crucial to overcome cleared environments while foraging in the anthropogenic matrix.

Our study evaluates, for the first time the scale of effect that best predicts phenotypic changes of a phyllostomid species in an anthropogenic landscape. Yet, incorporating the multivariate nature of wing morphology can bring more detailed and valuable insights to understand how flight-related ecomorphological traits of bats are shaped by landscape filtering—for example with morphometric geometry, see De Oliveira et al. (2020)—. Evidence from other phyllostomid species indicates that landscape transformation may impact the genetic diversity of populations depending on their movement capabilities (Meyer et al. 2009). This type of information joined with morphological data might shed light on how the response of *A. planirostris* is driven by natural phenotypic plasticity or, by contrast, is the result of microevolutionary processes associated with landscape anthropization (Alberti et al. 2017). Further studies incorporating morphology and bat trophic interactions will undoubtedly broaden our understanding of the effects of morphological inter-individual variation with the species ecosystem function.

## Acknowledgments

Natacha Chacoff and Tobías Rojas made helpful comments that improved the manuscript. We would like to thank Irene M. A. Bender for her constructive revision of the paper. We had discussions with Lilia Roa and María Ángela Echeverry-Galvis that enriched the project’s conceptual development. We are grateful to the members of the Grupo de Mamíferos Silvestres - Unillanos for their field assistance. During the preparation of the paper, Andrés F. Ramírez-Mejía was funded by a Ph.D. fellowship awarded by the Consejo Nacional de Investigaciones Científicas y Técnicas de Argentina (CONICET).

## Authors contribution

FS acquired financial support. AFRM and FS conceived and conceptualized the initial idea. AFRM, NUC, and FS made the sampling design. AFRM and FS did the fieldwork. FS provided laboratory resources and field equipment to collect the data. AFRM, FS, and NUC analyzed the data. AFRM wrote the first draft; FS and NUC critically reviewed and edited the manuscript.

## Ethical statement

We worked under permit 1313, of October 16, 2016, by the Colombian National Authority of Environmental Licenses (ANLA).

