## Supplementary material for "The interplay of spatial scale and landscape transformation moderates the abundance and intraspecific variation in the ecomorphological traits of a phyllostomid bat": Suplementary Figures S1 - S6

### Suplementary material

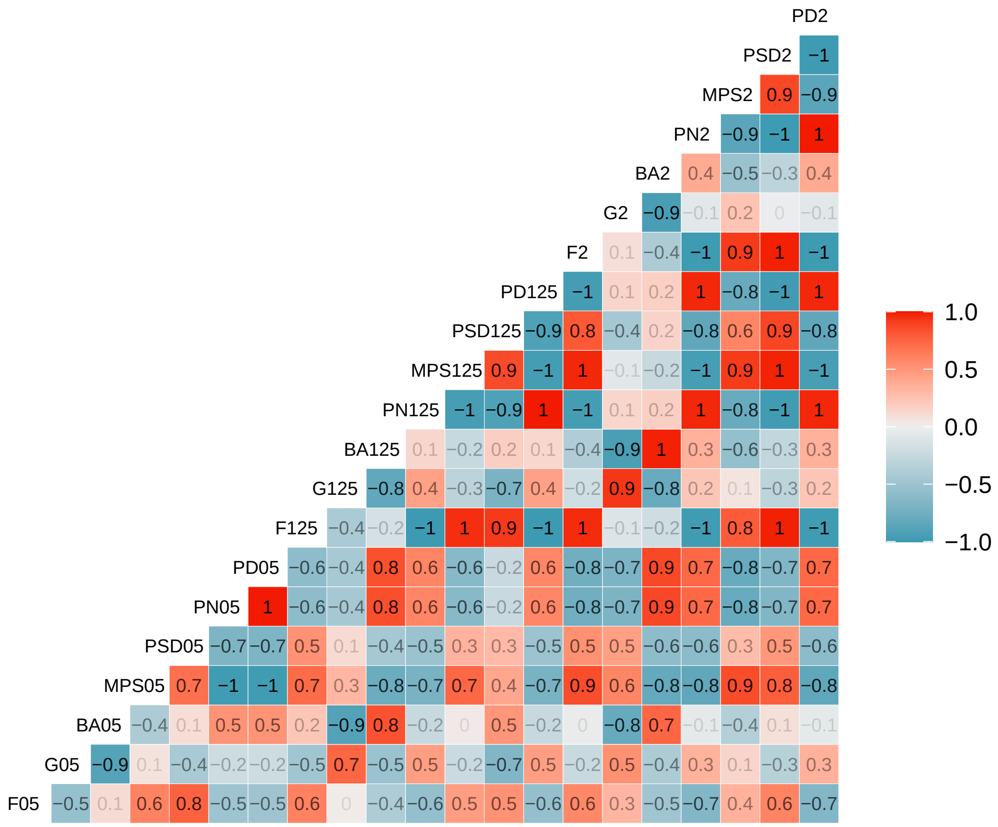


Figure S1. Pearson's correlation matrix of landscape predictor variables. BA = built area, F = forest, G = grassland, MPS = mean patch size, PD = patch density, PN = patch number, PSD = patch size standard deviation, 05 = 0.5 km scale, 125 = 1.25 km scale, 2 = 2 km scale.

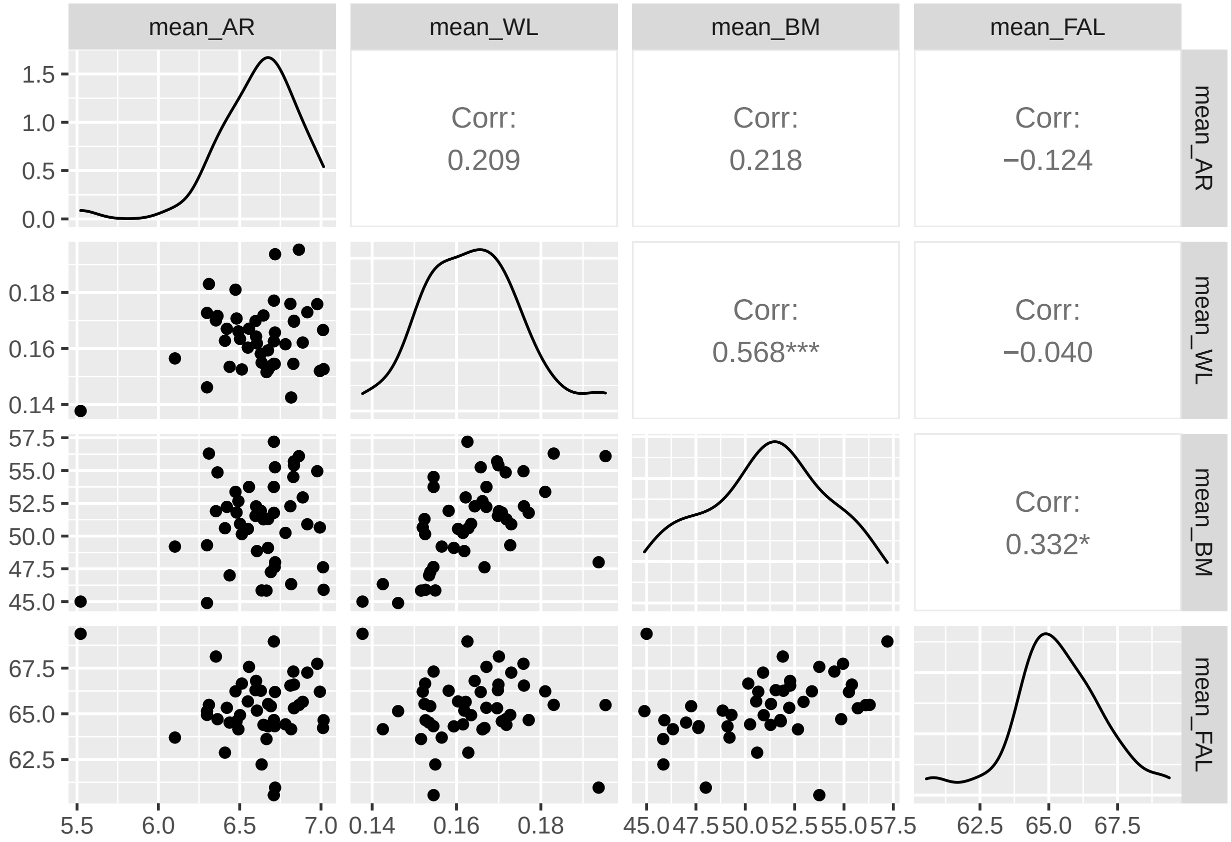


Figure S2. Pearson's correlations of *A. planirostris* ecomorphological traits. FAL = forearm length, AR = wing aspect ratio, WL = wing loading, BM = body mass. Diagonal density plots show the distribution of each variable in the population. One asterik indicates significant correlations (p<0.05), two asteriks indicate highly significant correlations (p<0.01).

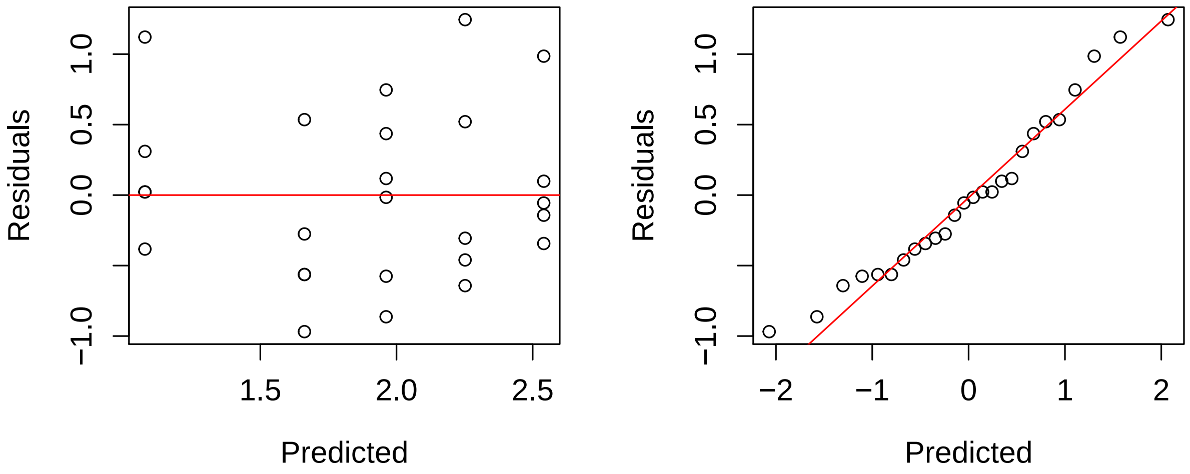


Figure S3. Residual analysis of best fitted model explaining changes of *A. planirostris* abundance as function of landscape variables. Left: residual vs. fitted values. Right: quantile-quantile plot of model's residuals.

This model included the grassland percentage at 2 km and 0.5 km, and the forest percentage ar 1.25 km.

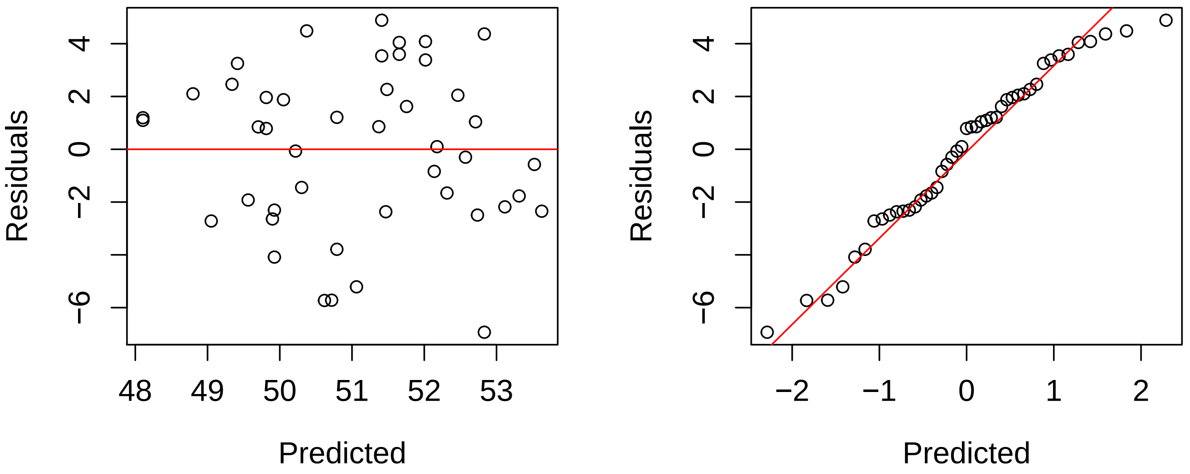


Figure S4. Residual analysis of best fitted model explaining changes of *A. planirostris* body mass (g) as function of landscape variables. Left: residual vs. fitted values. Right: quantile-quantile plot of model's residuals. This model included tha grassland percentage at 2 km and 0.5 km, and the forest percentage at 1.25 km and 0.5 km.

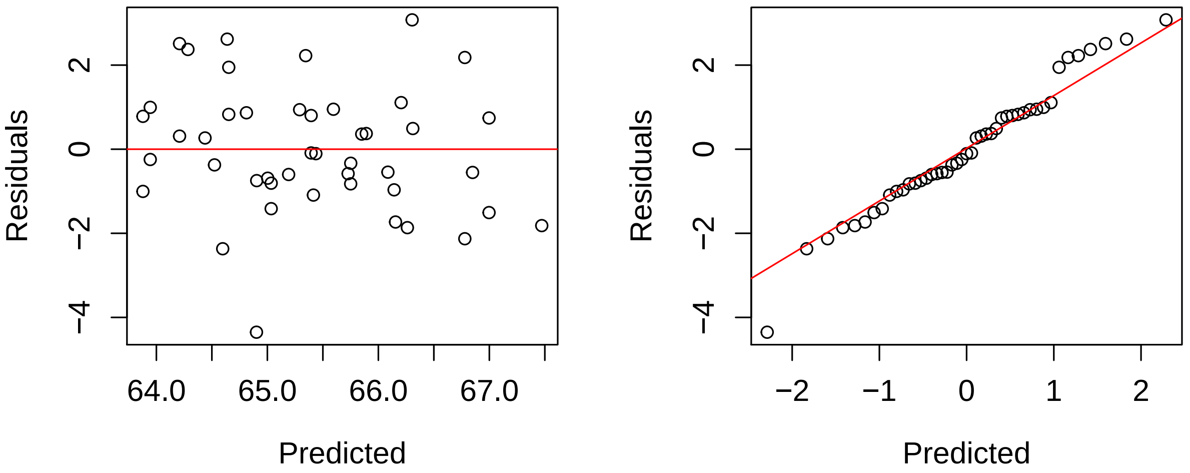


Figure S5. Residual analysis of best fitted model explaining changes of *A. planirostris* forearm length (mm) as function of landscape variables. Left: residual vs. fitted values. Right: quantile-quantile plot of model's residuals. This model included tha grassland percentage at 2 km and 0.5 km, and the forest percentage at 1.25 km and 0.5 km.

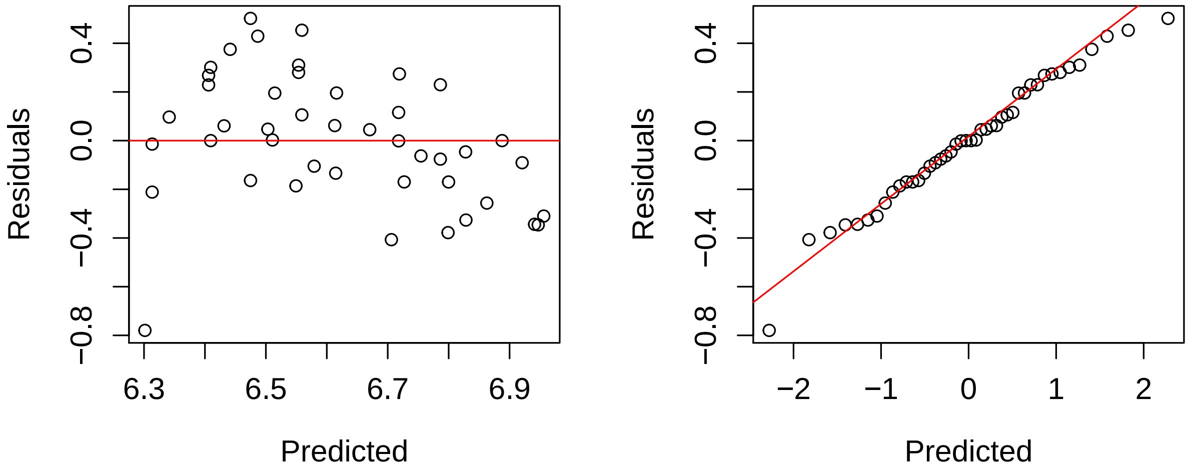


Figure S6. Residual analysis of best fitted model explaining changes of *A. planirostris* wing aspect ratio as function of landscape variables. Left: residual vs. fitted values. Right: quantile-quantile plot of model's residuals. This model included tha grassland percentage at 2 km and 0.5 km, and the forest percentage at 1.25 km and 0.5 km.
